# Extracellular vesicles derived from cells overexpressing HGSNAT rescue defects in Mucopolysaccharidosis IIIC neurons

**DOI:** 10.64898/2026.07.22.740105

**Authors:** Travis Moore, Mahsa Taherzadeh, Xuefang Pan, Melissa Hewitt, Caroline Sodja, Dylan Layton-Matthews, Mehrnaz Fazeli, Anahita Bakhshizadeh, Claudie Charlebois, Marina Rukhlova, Thomas Durcan, Anahita Bakhshizadeh Gashti, S. Mehdy Elahi, Jagdeep K. Sandhu, Anna Jezierski, Alexey V. Pshezhetsky

## Abstract

Mucopolysaccharidosis III type C (MPS IIIC) is a rare neurological lysosomal storage disorder caused by genetic deficiency of the lysosomal membrane enzyme, heparan-α-glucosaminide N-acetyltransferase (HGSNAT). To assess the feasibility of therapeutic strategies based on cross-correction of neurons by HGSNAT secreted from transplanted cells overexpressing the enzyme, we generated induced cortical neurons (iCN) from induced pluripotent stem cells (iPSCs) derived from MPS IIIC patients. The neurons were treated with extracellular vesicles (EV) purified from the culture medium conditioned by human endothelial cells transduced with a lentiviral vector encoding EGFP-tagged HGSNAT (LV-HGSNAT-EGFP). The isolated EV showed supraphysiologic HGSNAT activity levels and efficiently delivered the enzyme to the lysosomes of MPS IIIC iCN reducing lysosomal size and restoring normal synaptic protein levels. EV-mediated delivery of HGSNAT to neurons was further confirmed by the analysis of MPS IIIC iCN either co-cultured with iPSC-derived MPS IIIC microglia (iMGL) transduced with LV-HGSNAT-EGFP or treated with the iMGL conditioned medium. MPS IIIC iCN co-cultured with iMGL overexpressing HGSNAT achieved a complete phenotypic rescue, including normalization of lysosomal size, and the levels of heparan sulfate, G_M2_-ganglioside, synaptic proteins and brain-derived neurotropic factor. Treatment of MPS IIIC iCNs with conditioned medium led to a partial defects correction. Our findings reveal the translational potential of EV-mediated enzyme delivery in MPS IIIC patients treated with LV-mediated haematopoietic progenitor stem cell gene therapy.

## Introduction

Mucopolysaccharidosis type III (MPS III or Sanfilippo disease) is one of the approximately 70 genetic metabolic conditions affecting the lysosomal function and collectively known as lysosomal storage disorders (LSD) [1]. MPS III accounts for approximately 10% of all LSD [2], and has the highest reported prevalence of 1 in 24,000 births in the Netherlands [3, 4]. Patients primarily exhibit the central nervous system (CNS) manifestations, with relatively mild somatic features [5]. The disease typically has an onset in early childhood (2–4 years) with behavioral abnormalities, followed by developmental delay and progressive cognitive decline. This ultimately leads to childhood dementia, severe disability and premature death, generally before the third decade of life [6, 7].

MPS III comprises of four genetic subtypes (A, B, C, and D), each caused by a deficiency of a distinct lysosomal enzyme required for catabolism of a glycosaminoglycan heparan sulfate (HS) [8, 9]. Specifically, MPS III type C (MPS IIIC) results from pathogenic variants in the gene encoding heparan-α-glucosaminide N-acetyltransferase (HGSNAT), an enzyme that catalyses the transmembrane acetylation of HS necessary for its intra-lysosomal degradation [10].

Currently approved therapeutic strategies for other neurological LSD include enzyme replacement therapy (ERT) either with direct CNS delivery or using the brain-penetrant enzymes, haematopoietic stem and progenitor cell (HSPC) transplantation, HSPC gene therapy, and substrate reduction therapy (SRT), reviewed in [11]. However, none of these approaches have yet received regulatory approval for MPS III [12]. A major barrier to therapeutic efficacy remains the blood–brain barrier (BBB), which restricts the widespread delivery of systemically delivered therapeutics to the CNS [13].

Recent treatment strategies for Sanfilippo disease have focused on direct intracranial or intrathecal delivery of HS-degrading enzymes or AAV9 vectors encoding them. In murine and canine models, these approaches reduced HS in the blood and cerebrospinal fluid and improved neurological outcomes [14–16]. However, human clinical trials have shown only modest effects with disease stabilization rather than reversal of CNS pathology [17, 18]. Furthermore, direct CNS delivery methods are invasive and carry procedural risks and complications.

The first gene-modifying treatment clinically approved for a neurological LSD was the HSPC lentiviral (LV) gene therapy for metachromatic leukodystrophy (MLD), caused by the genetic deficiency of arylsulfatase A (*ARSA*). In this approach, patient’s HSPC are transduced *ex vivo* with a LV vector encoding *ARSA*, and subsequently transplanted. This leads to a sustained enzyme overexpression and secretion in all HSPC-derived cell lineages [19]. Notably, HSPC-derived macrophages migrate into the brain parenchyma and secrete ARSA to cross-correct the neighboring neurons. Treated patients have shown long-term normalization of ARSA activity in the periphery and CSF, halted disease progression, and sustained neurological benefits at 7–24 months after treatment [19, 20]. Those treated before symptom onset remained disease-free for more than 10 years [21].

A potential key challenge in implementing the HSPC LV gene therapy for MPS IIIC is related to the fact that the HGSNAT enzyme is a multi-spanning transmembrane protein, uncapable of free diffusion between the cells. One potentially promising but underexplored strategy to overcome this complication is the use of extracellular vesicles (EV), small, naturally secreted, membrane-bound particles capable of transferring functional proteins between cells, as vehicles to deliver therapeutic proteins incorporated in their membrane [22, 23]. In this study, we evaluated an HGSNAT delivery strategy using the enzyme-enriched EV (HGSNAT^+^ EV) secreted by gene-modified cells producing supraphysiological levels of the enzyme [24, 25]. We studied the potential of purified HGSNAT^+^ EV to correct pathophysiological features of cortical neurons (iCN) derived from MPS IIIC induced pluripotent stem cells (iPSCs). We also used MPS IIIC iPSC-derived microglia (iMGL) transduced with the HGSNAT-encoding LV to assess their ability to cross-correct cocultured MPS IIIC iCN [26, 27].

Our results demonstrate a robust normalization of disease biomarkers in MPS IIIC iCN treated with purified HGSNAT^+^ EVs or co-cultured with gene-corrected MPS IIIC iMGL. These findings suggest that the approaches based on therapeutic delivery of enzyme-loaded EV or HSPC LV gene therapy represent viable strategies for the treatment of MPS IIIC and other neurological LSD caused by genetic defects affecting lysosomal membrane enzymes and proteins.

## Results

### 1. Extracellular vesicles secreted by cultured hCMEC/D3 cells overexpressing HGSNAT-EGFP deliver the enzyme for cross-correction of other cells

Previously HGSNAT was detected in the EV secreted by primary cells, but its content was sparce reflecting low abundance of the enzyme in most of tissues [28]. Since the content of lysosomal enzymes is, typically, greatly increased in the EV secreted by cells overexpressing them, we transduced immortalized human cerebral microvascular endothelial cells (hCMEC/D3) with LV encoding the wild-type (WT) human EGFP-tagged HGSNAT, under the control of the CMV promoter (CMV-HGSNAT-EGFP LV) [25]. We chose hCMEC/D3 cells, which are of brain endothelial origin [29], to generate EVs with the lipid, protein, and glycan compositions resembling the microvasculature endothelium, thereby potentially enhancing their blood-brain barrier (BBB) tropism and transcytosis [30]. Forty-eight h after transduction, cells stably expressing EGFP were isolated by cell-sorting and propagated. The transduced hCMEC/D3 cells expressed EGFP, in a perinuclear punctate pattern typical of lysosomal proteins. Furthermore, EGFP fluorescence signal colocalized with the lysosomal marker, LysoTracker Red (**Figure S1B**). HGSNAT activity in stably transduced hCMEC/D3 cells increased ∼100-fold compared to the endogenous levels in non-transduced cells (**Figure S1C**).

EV were isolated from the medium conditioned by transduced hCMEC/D3 cells collected after 24 h using differential ultracentrifugation, following previously published protocols [31]. The isolated EV were characterized by nanoparticle tracking analysis using a ZetaView instrument, to determine their size, and concentration. The hCMEC/D3-derived EV had a mean diameter of ∼200 nm and were present at concentrations ranging from 1.5 x 10^11^ to 4.3 x 10^11^ EV/ml of medium (**Figure 1A**). The presence of the HGSNAT-EGFP fusion protein in the purified EV was confirmed by immunoblotting, which detected the presence of processed lysosomal HGSNAT 20 kDa protein chain, along with the precursor and oligomer forms of the enzyme (**Figure 1B**). Western blot analysis using Capillary immunoblot electrophoresis (ProteinSimple Western system) demonstrated the presence of pan-exosome markers such as CD63 and flotillin-2, and the absence of an endoplasmic reticulum (ER) marker calnexin, in the isolated EV (**Figure 1C**). These results suggest that the EVs primarily originated from late endosomes or multivesicular bodies.

**Figure 1:**
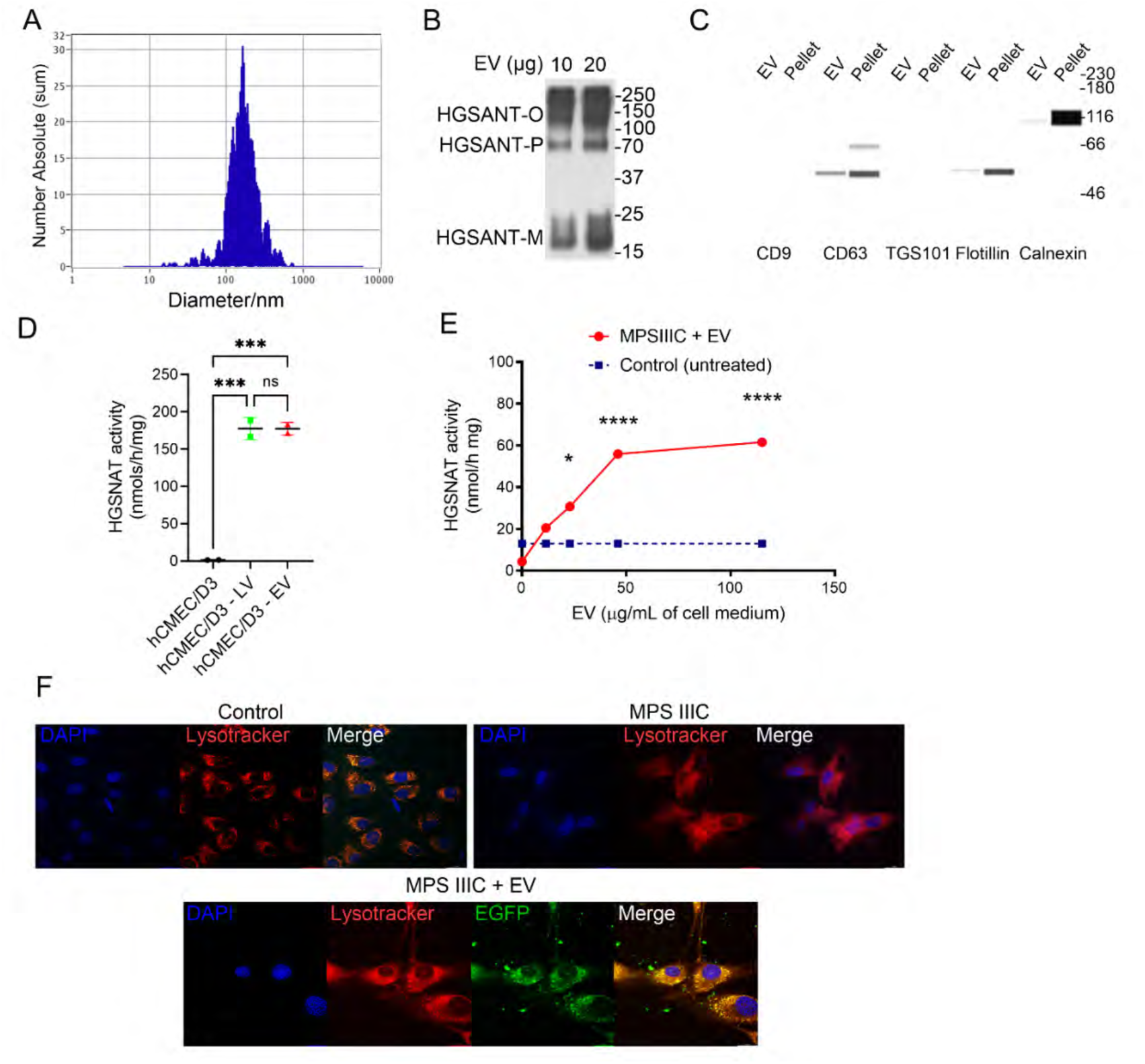
hCMEC/D3-derived EV contain exosome protein markers and HGSNAT-EGFP and deliver supraphysiologic levels of HGSNAT-GFP to the lysosomes of cultured fibroblasts of MPS IIIC patient. (**A**) EV isolated from the culture medium conditioned by hCMEC/D3 overexpressing HGSNAT-EGFP were characterized by the nanoparticle tracking analysis using Zetaview®. EV had an average size of ∼200 nm, characteristic for exosomes, and were present at a concentration of 1.5 - 4.3 x 10^11^ EV/ml of the medium. (**B**) Immunoblot analysis of EV at 10 and 20 µg/lane showed the presence of HGSNAT in the form of oligomers (O), precursor (P) and the mature form of the enzyme (M) found in the lysosomes. (**C**) Capillary immunoblot showed the presence of the EV-specific markers such as CD63 and Flotillin in EVs isolated from hCMEC/D3 cells. EVs were negative for the endoplasmic reticulum (ER) maker, calnexin, showing the absence of cellular contamination. (**D**) HGSNAT activity in the isolated EV is similar to that in post-sorted LV CMV-HGSNAT-EGFP transfected hCMEC/D3 cells and is >100 fold higher that the endogenous level in non-transduced hCMEC/D3 cells. (**E**) MPS IIIC fibroblasts treated with HGSNAT^+^ EV (11-115 mg/mL of cell medium for 18 h) show levels of HGSNAT activity exceeding the endogenous level in untreated control cells. (**F**) Representative IF images of human fibroblasts of a healthy control, MPS IIIC patient and of a MPS IIIC patient treated with isolated HGSNAT^+^ EV (46 µg/ml for 18 h). MPS IIIC cells treated with HGSNAT^+^ EV show presence of EGFP^+^ puncta in the cytoplasm colocalizing with lysosomal marker, Lysotracker red. DAPI (blue) was used to detect nuclei. Scale bars equal 25 µm.

HGSNAT enzymatic activity measured in the collected EV was comparable to that in the pelleted LV-transduced hCMEC/D3 cells (177.6±14.9 nmol/mg h) (**Figure 1D**). We further tested whether the HGSNAT^+^ EV could deliver the HGSNAT-EGFP enzyme to lysosomes of other cells, including MPS IIIC patient fibroblasts and non-transduced hCMEC/D3 cells. In brief, isolated HGSNAT^+^ EV were added to cultured MPS IIIC fibroblasts in concentrations ranging from 10 and 115 µg/mL of culture medium. After 18 h, cells were either harvested to measure HGSNAT activity or labeled with LysoTracker red and analyzed by confocal microscopy. EV-treated MPS IIIC fibroblasts revealed a marked increase in HGSNAT activity, reaching levels similar or exceeding those in healthy control cells. Confocal imaging revealed EGFP^+^ lysosomes, consistent with lysosomal uptake of HGSNAT-EGFP (**Figure 1E, F**). Additionally, treated cells displayed somewhat reduced LysoTracker red staining, indicative of decreased lysosomal storage. Similarly, hCMEC/D3 cells treated with HGSNAT-EGFP^+^ EV for 18 h (**Figure 1F**) demonstrated the presence of EGFP in the Lysotracker^+^ lysosomes, confirming that EV derived from HGSNAT-overexpressing cells can effectively deliver HGSNAT cargo to the lysosomes of target cells.

### 2. Treatment of MPS IIIC iPSC-derived cortical neurons with HGSNAT^+^ EV reduces the lysosomal storage and rescues deficits of synaptic proteins

We further evaluated whether the isolated HGSNAT^+^ EV secreted by hCMEC/D3 cells could rescue the lysosomal storage phenotype in iCN derived from MPS IIIC patient iPSCs and restore previously reported neuronal defects, such as decreased levels of synaptic puncta [32]. EV were added to cultured mature iCN at day in vitro (DIV) 25 at a final concentration of 21 µg/ml of culture medium (1.3 x 10^8^ EV/ml), which was found to be sufficient to increase HGSNAT activity in the recipient cells to or above the WT level (**Figure 1**). At the point of maturation (DIV 28), the cells were fixed and analyzed by immunofluorescent (IF) microscopy (**Figure 2).** This analysis revealed that EV-treated iCN were positive for HGSNAT-EGFP, which co-localized with the lysosomal marker LAMP2 (**Figure 2A**). Treated iCN also exhibited a 10-fold reduction in LAMP2^+^ areas compared to untreated iCNs, reaching levels similar to those observed in two lines of healthy control neurons (CNT-1 and CNT-2) (**Figure 2B**). These results suggest that EVs containing the HGSNAT enzyme effectively rescue the lysosomal storage phenotype in MPS IIIC iCNs.

**Figure 2:**
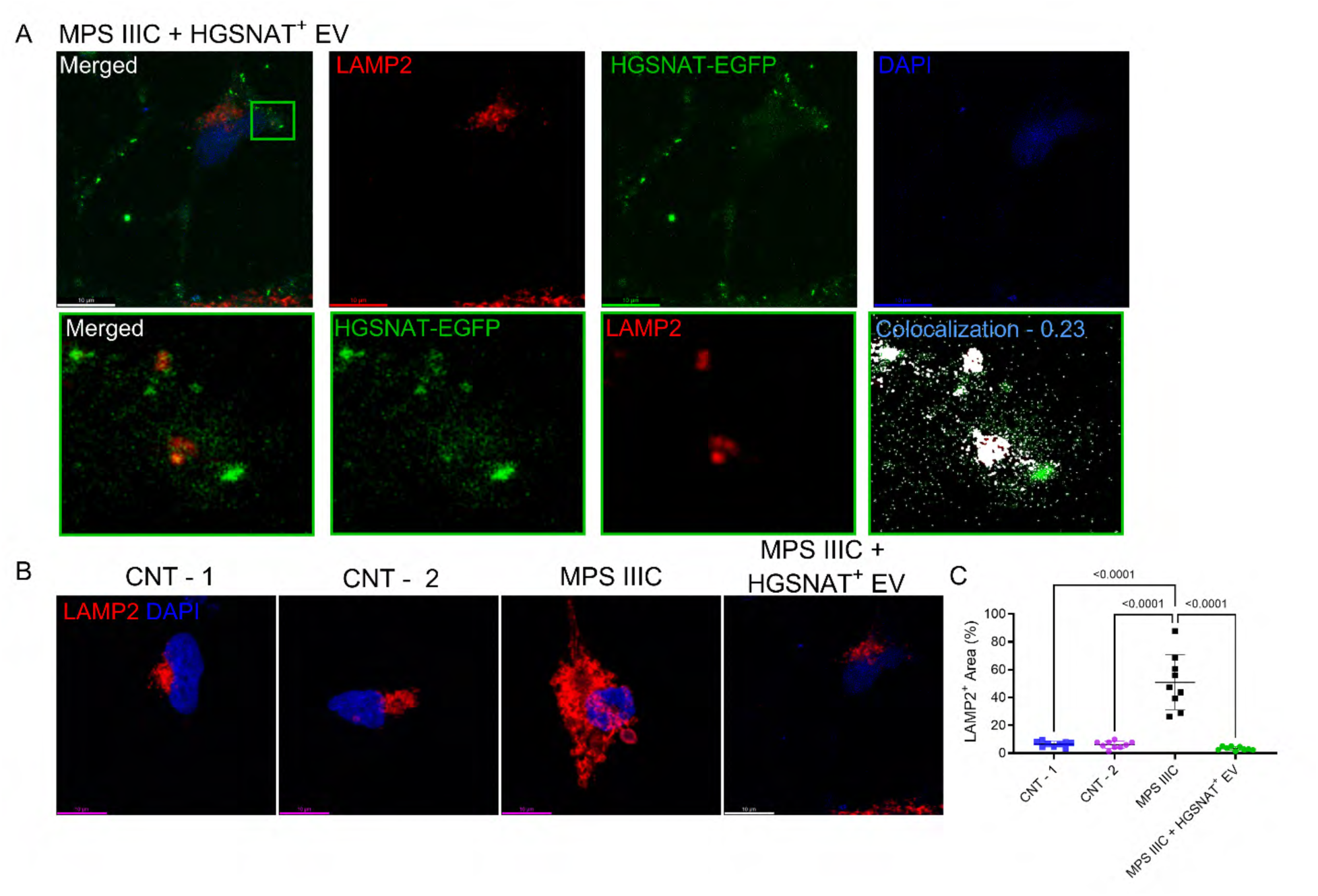
MPS IIIC iCN treated with HGSNAT^+^ EV show presence of HGSNAT-GFP in the lysosomes and rescue of the lysosomal storage phenotype. (**A**) Representative IF images of MPS IIIC iCN treated with HGSNAT^+^ EV and labeled for the lysosomal marker LAMP2 (red), and HGSNAT-EGFP (green). Zoomed images (lower panels) of the boxed area in the top images illustrate colocalization of LAMP2^+^ and EGFP^+^ puncta. DAPI (blue) was used to detect nuclei. Colocalization pixel map (white indicates areas of high colocalization) of the zoomed regions are shown at the bottom right. Person’s colocalization coefficient (r = 0.23) was calculated using the ImageJ colocalization threshold tool. (**B**) Representative images of iCN labeled for the lysosomal marker, LAMP2 (red), and DAPI (blue) (**C**) Quantification of LAMP2^+^ area (% of the total cell area) measured using ImageJ software. The LAMP2^+^ area in HGSNAT^+^ EV-treated MPS IIIC iCN was comparable to that of two healthy control cell lines (CNT-1 and CNT-2) and significantly reduced compared to untreated MPS IIIC cells. Data represent individual values, means and standard deviations (SD) from 3 independent experiments are shown (>3 cells per biological replicate). *P*-values were calculated by nested one-way ANOVA and Tukey’s post hoc test. Scale bars equal 10 µm.

The MPS IIIC iCN treated with HGSNAT^+^ EV were also assessed for synaptic integrity by quantifying puncta positive for makers of glutamatergic (VGLUT1 and PSD95) and GABAergic (vGAT and Gephyrin) synapses, as well as for brain-derived neurotrophic factor (BDNF), which we previously found to be reduced in MPS IIIC iCN [25, 33] (Moore., Dubot., et al., bioRxiv 2026.02.20.707013). MPS IIIC iCN treated with HGSNAT^+^ EVs showed a pronounced increase in the numbers of VGLUT1^+^, VGAT^+^ and BDNF^+^ puncta compared to untreated MPS IIIC iCNs to the levels similar to those observed in healthy controls iCN (**Figure 3A-C**). In contrast, the number of puncta positive for the excitatory post-synaptic marker, PSD95, did not significantly change following EV treatment. The number of puncta positive for the inhibitory post-synaptic marker, Gephyrin, increased significantly in treated compared to untreated MPS IIIC iCN but remained below the control levels. It’s tempting to speculate that a longer treatment duration and/or multiple applications of HGSNAT^+^ EV may be required for delivering a therapeutically relevant dose to achieve a full recovery of post-synaptic neuronal function. Overall, these results suggest that EV carrying the HGSNAT enzyme, restore disease-associated phenotypes in mature MPS IIIC iCN, improving both lysosomal homeostasis and synaptic integrity.

**Figure 3.**
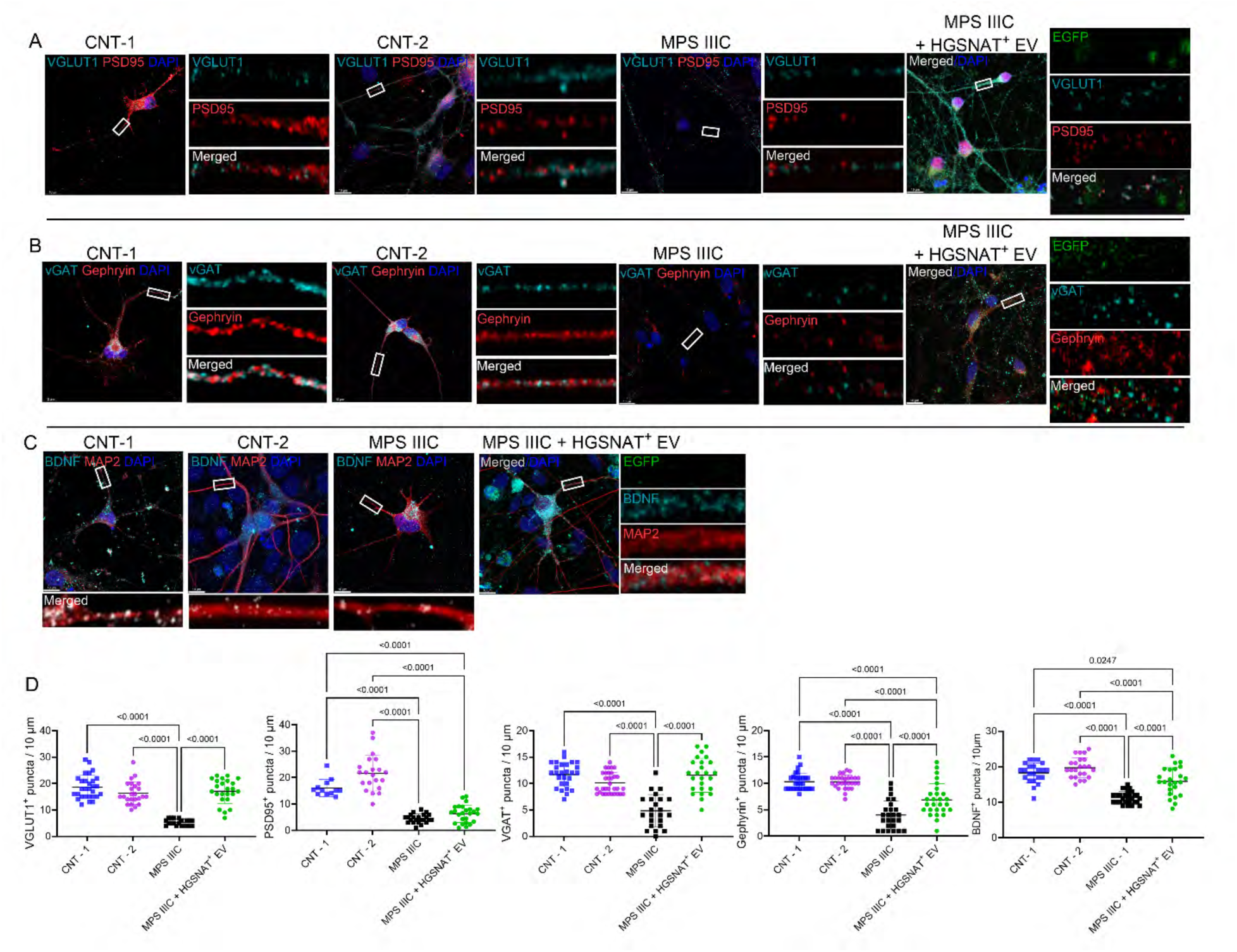
MPS IIIC iCN treated with HGSNAT^+^ EV show rescue of deficient levels of pre-synaptic protein markers VGLUT1 and VGAT and increased levels of Gephyrin and BDNF. **(A)** Deficient levels of the excitatory synapse marker, VGLUT1^+^ puncta in MPS IIIC iCN are rescued by HGSNAT^+^ EV treatment. Panels show representative images of healthy control (CNT-1, and CNT-2) and MPS IIIC neurons treated or not with HGSNAT^+^ EV immunolabeled for EGFP (green), VGLUT1 (cyan) and PSD95 (red). **(B)** Deficient levels of protein markers of the inhibitory synapse, vGAT^+^ and Gephyrin^+^ puncta in MPS IIIC iCN are rescued by HGSNAT^+^ EV treatment. Panels show representative images of healthy control (CNT-1, and CNT-2) and MPS IIIC neurons treated or not with HGSNAT^+^ EV and labeled for EGFP (green), VGAT (cyan) and Gephyrin (red). **(C)** Deficient levels of BDNF^+^ puncta in MPS IIIC iCN are increased by HGSNAT^+^ EV treatment. Panels show representative images of MPS IIIC neurons and those of healthy controls (CNT-1, and CNT-2) treated or not with HGSNAT^+^ EV and labeled for EGFP (green), BDNF (red) and neuronal dendrite marker MAP2 (grey). In all panels, DAPI (blue) was used to label nuclei. The scale bars equal 10 µm. Inserts show enlarged images of neurites selected at ≥10 µm from the soma (white rectangles). The graphs show counts of VGLUT1^+^, PSD95^+^, VGAT^+^, Gephyrin^+^, and BDNF^+^ puncta along 10-µm segments of neuronal projections by ImageJ software. Individual values, means and SD from 3 biological replicates (>15 cells in each experiment) are shown. *P*-values were calculated using nested one-way ANOVA and Tukey post hoc test.

### 3. iPSC-derived MPS IIIC microglia aggravate defects in co-cultured MPS IIIC iCN

To determine whether HGSNAT^+^ EV are also secreted by cultured microglia overexpressing the enzyme, we generated induced microglia (iMGL) from iPSC of MPS IIIC patients and healthy controls using recently developed differentiation protocols [26]. The iMGL phenotypes were evaluated in both monocultures and in co-cultures with their respective neuronal lines, i.e., control iCN with control iMGL, and MPS IIIC iCN with MPS IIIC iMGL. We also tested whether iCN co-cultured with iMGL had altered neuronal phenotypes compared to iCN monocultures.

iMGL monocultures were generated and matured to DIV 25, plated onto Matrigel^TM^-coated coverslips, and further matured to DIV 28. Cells were then fixed and immunolabelled for the pan-microglia marker IBA1, and the lysosomal marker, LAMP2 (**Figure 4A**) or for LAMP2 and HS (**Figure 4B**). Similarly to the monocultures of MPS IIIC iCN, MPS IIIC iMGL showed increased HS^+^ and LAMP2^+^ areas, compared to their respective controls (**Figure 4A-C**). The cells were also stained with fluorescently-labeled isolectin B4 (ILB4), which shows an increased affinity for activated microglia [34]. MPS IIIC iMGL showed an increased ILB4 labeling compared to control cells, indicative of a pro-inflammatory phenotype (**Figure 4A**).

**Figure 4:**
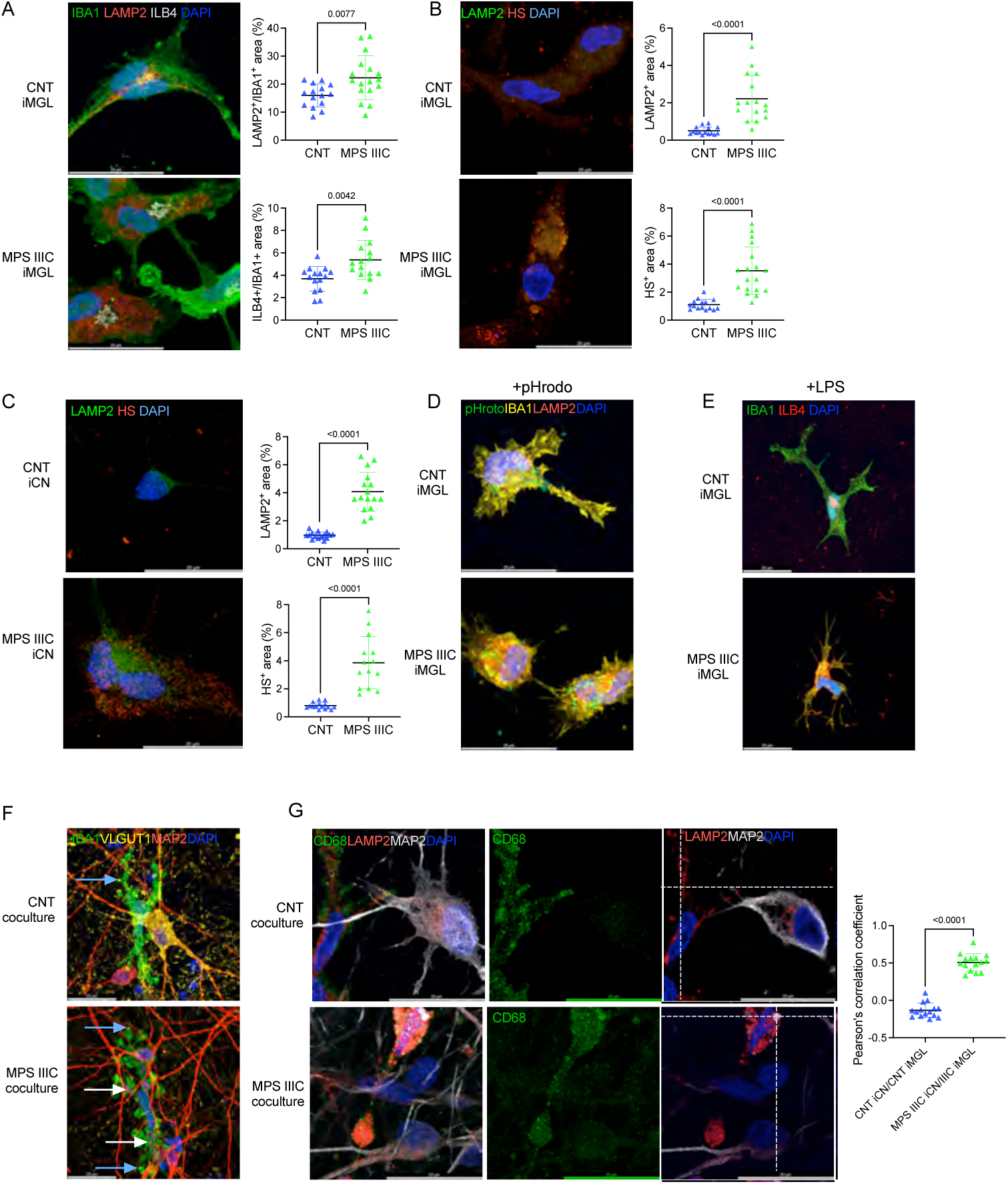
Biomarkers of MPS IIIC iMGL monocultures and iMGL/iCN co-cultures. (A-E) Control CNT and MPS IIIC iMGL in monocultures express characteristic microglia markers, phagocytic competence and respond to inflammatory stimuli, while the MPS IIIC iMGL have a distinct activated and lysosomal storage phenotype. **(A)** Representative images of healthy control (CNT) and MPS IIIC patient DIV 28 iMGL labeled for the microglial marker IBA1 (green), lysosomal marker LAMP2 (red), and the marker of activated microglia ILB4 (grey). Quantification with ImageJ software shows an increase of both LAMP2^+^ and ILB4^+^ areas in MPS IIIC compared to CNT iMGL. **(B-C)** Representative images for CNT and MPS IIIC iMGL **(B)** and iCN **(C)** labeled for LAMP2 (green) and HS (red). Quantification with ImageJ software shows an increase of LAMP2^+^ and HS^+^ areas in both MPS IIIC iMGL and iCN compared to their respective control cells. **(D)** Representative images for CNT and MPS IIIC iMGL exposed to pHrodo™ bacteria (green) for 1 h and labeled for LAMP2 (red), and IBA1 (yellow). Both MPS IIIC and CNT cells show phagocytic competence. **(E)** Representative images for CNT and MPS IIIC iMGL exposed to 100 ng/ml LPS for 24 hrs and labeled for IBA1 (green), BDNF (red), and ILB4 (grey). Cells show ameba-like shape suggesting their responsiveness to pro-inflammatory stimuli. (**F**) Representative images of DIV 35 CNT and MPS IIIC iMGL/iCN co-cultures. The cells were labeled for the dendrite marker MAP2 (red), microglia marker IBA1 (green), and pre-synaptic neuronal marker VGLUT1 (yellow). Both CNT and MPS IIIC iMGL contain VGLUT1^+^ puncta (blue arrows) while only MPS IIIC iMGL contain MAP2^+^ puncta (white arrows) consistent with active scavenging of neuronal processes. (**G, left and central panels**) Representative images DIV 35 co-cultures of CNT and MPS IIIC iMGL/iCN labeled for LAMP2 (red), MAP2 (grey), and the activated microglia marker CD68 (green). (**G, right panels**) XY sections from a single Z-stack for the images on the left, showing a colocalization of MAP2^+^ (grey) and LAMP2^+^ (red) areas at the dotted lines crosshairs in MPS IIIC iMGL not present in CNT iMGL. Colocalization threshold map displays colocalization of MAP2^+^ (white) and LAMP2^+^ (red) areas. Co-localization thresholds quantification of MAP2^+^ puncta within LAMP2^+^ areas of iMGL with ImageJ software shows colocalization in MSP IIIC but not in CNT iMGL. All graphs show individual results, means and s.e.m. from three independent cultures (5 images per culture). *P*-values were calculated by unpaired *t*-test with Welch’s correction. In all panels DAPI (blue) was used to label nuclei. The scale bars equal 25 µm.

We further treated control and MPS IIIC iMGL with pHrodo™ particles and *Escherichia coli* liposaccharide (LPS, 100 ng/ml) [27] to assess their phagocytic capacity and responsiveness to pro-inflammatory stimuli, respectively (**Figure 4D,E**). Both control and MPS IIIC iMGL internalized pHrodo™ (**Figure 4D**) and acquired an activated amoeboid morphology in response to LPS stimulation (**Figure 4E**), indicating functionally mature microglia phenotype. Neuronal/microglial co-cultures were then established by replating DIV 28 iMGL onto DIV 28 iCN grown on Poly-ornithine/laminin-coated glass coverslips, followed by an additional seven days of co-culture in a 1:1 mixture of iCN and iMGL media. At DIV 35, the cells were fixed and analyzed by IF to assess the levels and localization of neuronal synaptic marker VGLUT1, pan-lysosomal marker LAMP2, neuronal projection marker MAP2, and the microglial markers IBA1 and CD68 (**Figure 4F,G**). As in monocultures, MPS IIIC iMGL showed increased LAMP2 levels in microglial/neuronal cocultures consistent with lysosomal storage. In co-cultures of MPS IIIC microglia with MPS IIIC neurons, we also observed VGLUT1^+^ puncta and MAP2^+^ puncta within the microglia cell bodies defined by IBA1^+^ areas (**Figure 4F**, blue and white arrows, respectively). Furthermore, in MPS IIIC co-cultures, MAP2^+^ materials was present inside the LAMP2^+^ microglia lysosomes, suggesting that activated MPS IIIC iMGL were actively engulfing and degrading neuronal processes and synaptic components (**Figure 4G**). This was consistent with drastically reduced levels of vGLUT1^+^ (**Figure 4F)** puncta and BDNF puncta (**Figure S2**) in MPS IIIC neuronal/microglial co-cultures. Quantitative analysis of colocalization between MAP2 and LAMP2 signals in CD68^+^ microglia, revealed significantly higher colocalization coefficients in MPS IIIC co-cultures than in control co-cultures (**Figure 4G**). These findings are consistent with aberrantly increased synaptic pruning activity and/or impaired lysosomal degradation in MPS IIIC iMGL. They also suggest that activated microglia contribute to the disease pathogenesis and progression in the MPS IIIC neurons.

### 4. Gene-corrected MPS IIIC iMGL overexpressing HGSNAT-EGFP rescue the disease phenotype in co-cultured MPS IIIC iCN

We further tested whether gene-corrected MPS IIIC iMGL transduced with LV-HGSNAT-EGFP could rescue or ameliorate the disease phenotype in co-cultured MPS IIIC iCN. Lentiviral transduction was performed at the stage of induced hematopoietic progenitor cells (iHPC), an intermediate phase of iPSC-to-iMGL differentiation, due to their high proliferative capacity [26]. We used the LV-CD68-HGSNAT-EGFP vector, in which the HGSNAT expression is driven by the human CD68 promoter. This promoter was expected to be highly active in MPS IIIC iMGL, based on their activated status and high CD68 levels detected by IF microscopy. The iHPC were transduced at Multiplicity of Infection (MOI) 30 which in our previous experiments with primary mouse HSPC resulted in average transduction efficiency of ∼85% with ∼2 integrated viral vector copies per host cell genome (VCN 2.0) (T. Khayer, unpublished). The transduced MPS IIIC iHPC were further matured in monocultures until DIV 28 and then collected for measurement of HGSNAT enzyme activity in cell pellets or fixed for IF analysis for previously described markers of lysosomal storage (LAMP2, HS).

Non-transduced MPS IIIC iHPC showed almost undetectable HGSNAT activity, whereas LV-CD68-HGSNAT-EGFP-transduced iHPC exhibited, on average, ∼7-fold increase in HGSNAT activity compared to healthy control iHPC (**Figure S3A**). Similar results were observed in mature (DIV 28) iMGL, with HGSNAT activity below detection level in non-transduced MPS IIIC cells or those transduced with the control LV-CMV-EGFP (**Figure S3B**). In contrast, iMGL derived from iHPC transduced with LV-CD68-HGSNAT-EGFP showed supraphysiologic HGSNAT activity increased ∼10-fold compared to normal control iMGL (**Figure 3SB**).

At DIV 28, MPS IIIC iMGL transduced or not with LV-CD68-HGSNAT-EGFP or LV-CMV-EGFP were seeded over iCN cultures. Co-cultures were maintained for 7 days, until DIV 35, and then collected for measurement of HGSNAT enzyme activity in total cell pellets or fixed for IF analysis. Similar analyses were also performed for MPS IIIC or control iCN treated for 72 h with the media conditioned by monocultures of normal control iMGL, and LV-CD68-HGSNAT-EGFP-transduced MPS IIIC iMGL.

MPS IIIC iCN co-cultured with LV-CD68-HGSNAT-EGFP-transduced iMGL showed 8-10 fold reduction in the levels of LAMP2^+^, HS^+^ and G_M2_^+^ areas compared to MPS IIIC iCNs cocultured with untreated MPS IIIC iMGL. These levels were similar to those observed in control iCN/iMGL co-cultures, consistent with a rescue of primary and secondary lysosomal storage (**Figure 5A-C**). The neurons also displayed increased densities of BDNF^+^, VGLUT1^+^, PSD-95^+^, VGAT^+^, and gephyrin^+^ puncta, comparable to healthy control iCN (**Figure 5D-F**). HGSNAT enzymatic activity measured in homogenates of total cell pellets, and reflecting the combined activity in iCN and iMGL, was undetectable in the co-cultures of MPS IIIC iCN and untransfected or LV-CMV-GFP-transfected MPS IIIC iMGL. In the co-cultures of MPS IIIC iCN and LV-CD68-HGSNAT-EGFP-transfected MPS IIIC iMGL, HGSNAT activity was increased to supraphysiological levels, ∼10-fold higher as compared to control iCN/iMGL co-cultures (**Figure 5G**). In contrast, HGSNAT enzymatic activity in MPS IIIC iCN/control iMGL co-cultures although higher than MPS IIIC iCN/MPS IIIC iMGL co-cultures remained lower than in control iCN/iMGL co-cultures (**Figure 5G).**

**Figure 5:**
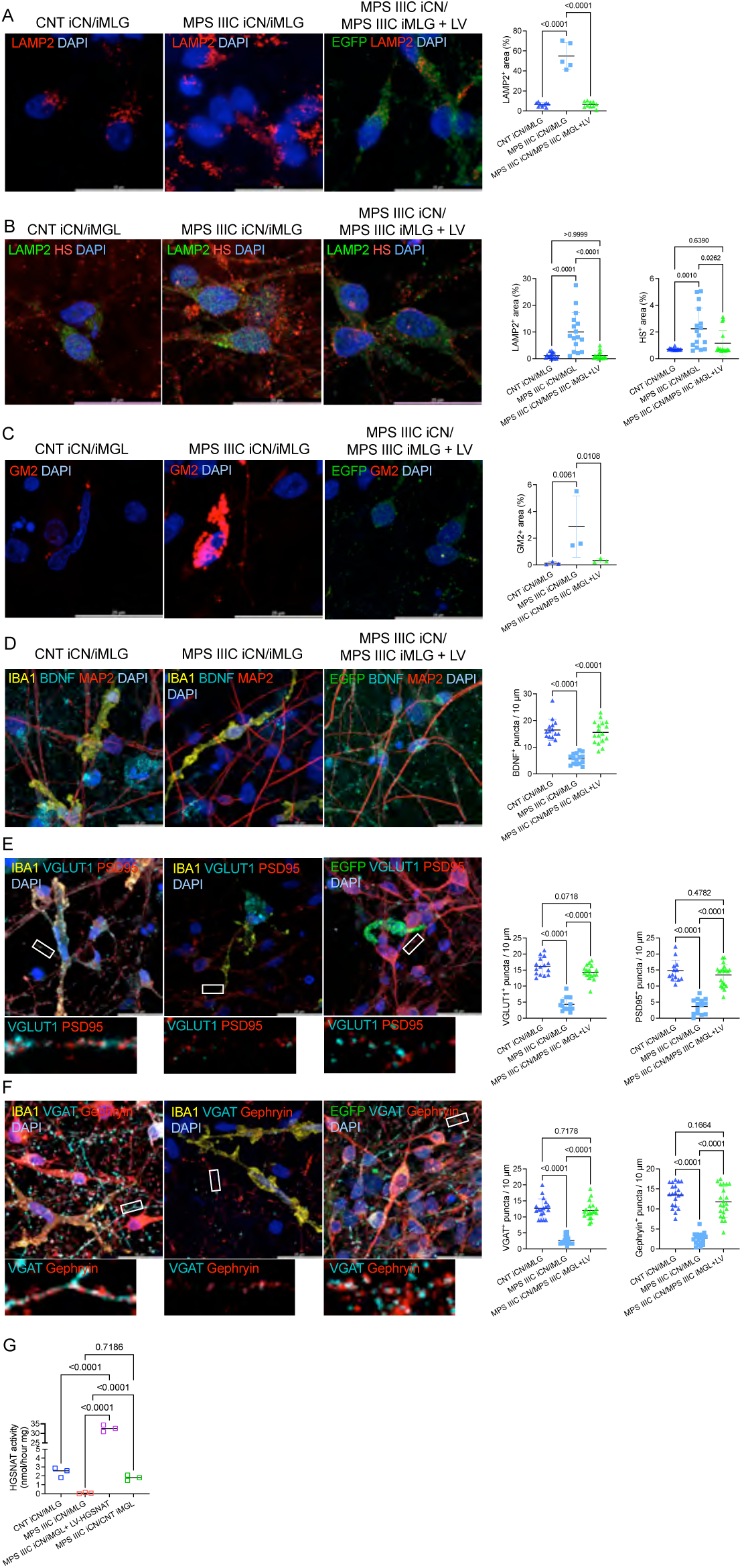
MPS IIIC iCN co-cultured with MPS IIIC iMGL transduced with LV CD68-HGSNAT-EGFP show rescue of lysosomal storage, secondary accumulation of G_M2_-ganglioside and deficient levels of BDNF and synaptic proteins. Panels show representative IF images of DIV 35 CNT and MPS IIIC iMGL/iCN co-cultures and co-cultures of MPS IIIC iCN with MPS IIIC iMGL transduced with the LV CD68-HGSNAT-EGFP vector. **(A)** Cells are labeled for LAMP2 (red) and HGSNAT-EGFP (green). The graph shows quantification of LAMP2^+^ area in iCN by ImageJ demonstrating reduction of LAMP2+ areas in MPS IIIC iCN co-cultured with iMGL overexpressing HGSNAT-EGFP to the level in control healthy cells. **(B)** Cells are labeled for LAMP2 (green) and HS (red). The graph show quantification of LAMP2^+^ and HS^+^ areas in iCN by ImageJ demonstrating reduction of LAMP2^+^ and HS^+^ areas in MPS IIIC iCN co-cultured with iMGL overexpressing HGSNAT-EGFP. **(C)** Cells are labeled for G_M2_ ganglioside (red) and HGSNAT-EGFP (green). Graph shows quantification of GM2^+^ areas with ImageJ software. Accumulation of GM2 is reduced in MPS IIIC iCN co-cultured with iMGL overexpressing HGSNAT-EGFP. **(D)** Cells are labeled for IBA1A (yellow), BDNF (cyan), MAP2 (red), and HGSNAT-GFP (green). The graphs show quantification of BDNF^+^ puncta along 10-µm segments of neuronal dendritic projections using ImageJ. Reduced levels of BDNF^+^ puncta are rescued in MPS IIIC iCN co-cultured with iMGL overexpressing HGSNAT-EGFP. **(E)** Cells are labeled for VGLUT1 (cyan) and PSD95 (red) and HGSNAT-GFP (green). Inserts show enlarged images of neurites selected at ≥10 µm from the soma (white rectangles). The graphs show quantification of VGLUT1^+^ or PSD95^+^ puncta along 10-µm segments of neuronal projections using ImageJ. Levels of VGLUT1^+^ and PSD95^+^ puncta are reduced in untreated MPS IIIC compared to CNT iCN and rescued in the MPS IIIC iCN cocultured with iMGL overexpressing HGSNAT-EGFP. **(F)** Cells are labeled for IBA1 (yellow), VGAT (cyan), Gephyrin (red) and HGSNAT-EGFP (green). Inserts show enlarged images of neurites selected at ≥10 µm from the soma (white rectangles). The graphs show quantification of VGAT^+^ or Gephyrin^+^ puncta along 10-µm segments of neuronal projections that are not overlayed by microglia with ImageJ. Levels of VGAT^+^ and Gephyrin^+^ puncta are reduced in MPS IIIC compared to CNT iCN and rescued in the MPS IIIC iCN cocultured with iMGL overexpressing HGSNAT-EGFP. (**G**) At DIV 35 MPS IIIC co-cultures show deficient HGSNAT activity. Co-cultures of MPS IIIC iCN and MPS IIIC iMGL transduced with LV CD68-HGSNAT-EGFP show ∼10-fold higher HGSNAT activity than co-cultures of control cells (CNT). In all panels DAPI (blue) was used to label nuclei. The scale bars equal 25 µm. Graphs show individual results, means and SD from three independent cultures (≥15 cells in each experiment). *P*-values were calculated using one-way ANOVA and Tukey post hoc test.

To determine whether the rescue of the deficits in MPS IIIC iCN cocultured with LV-CD68-HGSNAT-EGFP transduced MPS IIIC iMGL resulted from cross-correction mediated by HGSNAT^+^ EV secreted by microglia overexpressing HGSNAT, we treated MPS IIIC iMGL monocultures with conditioned media of either LV-transduced MPS IIIC iMGL or non-transduced control iMGL for 72 h (**Figure 6**). The collected media was centrifuged following the published protocol to remove large cellular debris and apoptotic bodies while retaining large EVs and exosomes [35]. The processed media was then added to fully differentiated control and MPS IIIC iCN in a 1:1 iMGL/iCN media ratio. Cells were subsequently maintained for 56 h before being analyzed by IF for lysosomal and synaptic markers as described above. MPS IIIC iCN treated with the conditioned media from LV CD68-HGSNAT-EGFP-transduced iMGL showed normalized LAMP2^+^ levels, and fully rescued levels of VGLUT1^+^, and BDNF^+^ puncta (**Figure 6A-D**). Similar to the MPS IIIC cultures treated with purified HGSNAT^+^ EV, no change was observed in PSD95^+^ puncta levels (**Figure 6D**). Interestingly, a partial reduction of LAMP2^+^ area was also detected in MPS IIIC iCN treated with the medium conditioned by control iMGL cultures (**Figure 6A**). However, other biomarkers (BDNF^+^, VGLUT1^+^, and PSD-95^+^ puncta) remained unchanged. This partial rescue suggests that the levels of HGSNAT^+^ EV in the medium conditioned by unmodified control iMGL is insufficient to achieve complete cross-correction of neurons.

**Figure 6:**
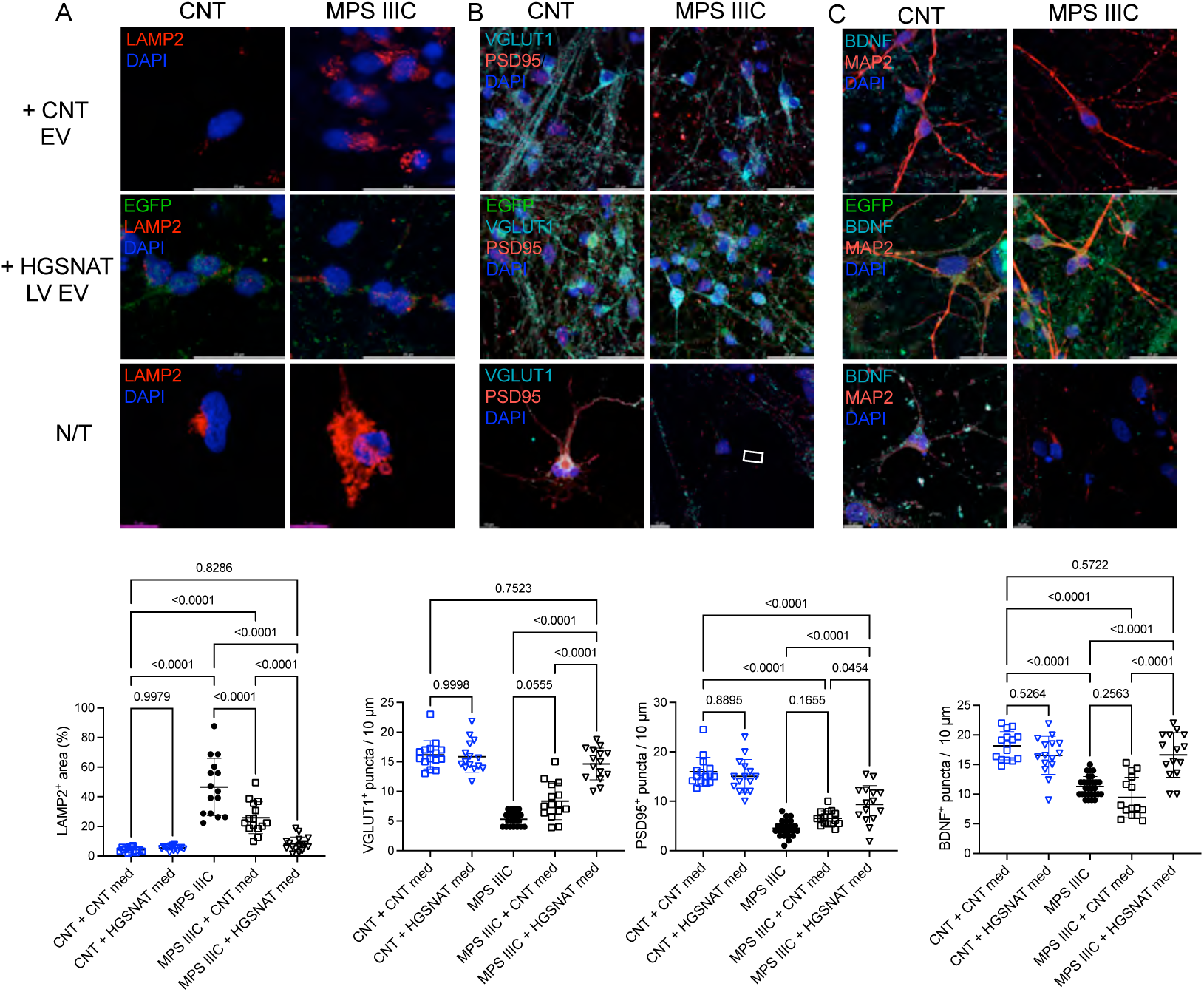
Treatment of MPS IIIC iCN with the media conditioned by MPS IIIC microglia overexpressing HGSNAT-EGFP leads to a partial rescue of lysosomal storage and synaptic deficits. Representative images from monocultures of control (CNT) or MPS IIIC iCN treated or not with the media conditioned either by CNT iMGL or MPS IIIC iMGL transduced with LV-CD68-HGSNAT-EGFP. **(A)** LAMP2^+^ area (red) is increased in untreated MPS IIIC iCN and normalised in the MPS IIIC iCN treated with the media conditioned by MPS IIIC microglia overexpressing HGSNAT-EGFP. In the cells treated with the media conditioned by CNT iMGL the LAMP2 levels are reduced but not normalized. **(B)** Reduced levels of VGLUT1^+^ puncta are rescued in the MPS IIIC iCN treated with the media conditioned by iMGL overexpressing HGSNAT-EGFP by not by the media conditioned by CNT iMGL. Deficient levels of PSD-95^+^ puncta are increased in MPS IIIC iCN treated with the media conditioned by MPS IIIC microglia overexpressing HGSNAT-EGFP but remain below the levels in CNT cells. **(C)** BDNF^+^ puncta are rescued in the MPS IIIC iCN treated with the media conditioned by iMGL overexpressing HGSNAT-EGFP by not by the media conditioned by CNT iMGL. The graphs show quantifications of LAMP2^+^ area (% of the total cell area) with ImageJ software or counts of puncta along 10-µm segments of neuronal dendritic projections. Individual results, means and SD from 3 biological replicates (≥5 cells/culture) are shown. *P*-values were calculated using nested one-way ANOVA and Tukey post hoc test.

## Discussion

Currently, there are no clinically approved therapies for treating the CNS manifestations of MPS III, underscoring the urgent need for novel treatment approaches. One promising avenue is LV-mediated *ex vivo* gene therapy, in which patient’s HSPC are gene-corrected and transplanted, resulting in sustained overexpression of the therapeutic enzyme in HSPC-derived cell lineages. CNS pathology is subsequently ameliorated as HSPC-derived microglia repopulate the brain and cross-correct resident cells such as neurons, astrocytes and oligodendrocytes (reviewed in [36]). This approach has shown efficacy for LSD caused by deficiencies in soluble lysosomal enzymes, such as arylsulfatase A in MLD or sulfaminidase in MPS IIIA. However, its suitability for MPS IIIC has been uncertain because the deficient enzyme, HGSNAT, is a transmembrane protein and does not readily diffuse between cells. Interestingly, previous work has shown that HGSNAT can be secreted from the cells as part of EV [28], which may enable delivery of the enzyme to target cells and potentially across the BBB. Here, using the human *in vitro* neuronal model, we sought to determine whether MPS IIIC patient-derived iCN can be rescued by delivery of HGSNAT-loaded EV, and whether induced microglia-like iMGL cells (iMGL) overexpressing HGSNAT could provide effective cross-correction of neighbouring neurons.

Our experiments established the feasibility of producing HGSNAT^+^ EV containing supraphysiologic levels of the enzyme, exceeding endogenous levels by more than 100-fold, from cultured hCMEC/D3 endothelial cells and iPSC-derived microglia transduced with HGSNAT-encoding LV vectors. When applied to cultures of skin fibroblasts and iPSC-derived iCN from MPS IIIC patients, the HGSNAT^+^ EV successfully delivered the enzyme to the lysosome compartment and reduced the lysosomal storage phenotype after a sustained treatment for only 18-54 h. Importantly, the key neuronal biomarkers, including the densities of BDNF^+^, VGLUT1^+^, and VGAT^+^ puncta, reduced in the MPS III cells approached normal control levels following treatment, indicating that tangible improvements can be achieved even in mature neurons. Conversely, post-synaptic markers (PSD-95^+^ and Gephyrin^+^ puncta) showed only partial restoration, suggesting stage-specific kinetics in lysosomal and postsynaptic marker recovery. Overall, our findings establish HGSNAT^+^ EV as a feasible and effective therapeutic vehicles for enzyme delivery and correction of lysosomal pathology and neuronal function in MPS IIIC.

Taking advantage of recently developed protocols for generating iPSC-derived iMGLs, we established and analysed co-cultures of MPS IIIC iCN with MPS IIIC iMGL, as well as with iMGL transduced using HGSNAT-encoding LV vectors and expressing ∼7-fold higher HGSNAT levels compared to the endogenous one. This system models post-LV-HSPC gene therapy brain environment, in which gene-corrected HSPC-macrophages overexpress HGSNAT and secrete it via EV, potentially providing a sustained source of the deficient enzyme for resident neurons. Importantly, the co-culture model enables bidirectional communication between MPS IIIC neurons and microglia, allowing investigation of how these cell types mutually influence disease phenotypes compared to their respective monocultures.

In MPS III, microglial activation triggered by HS fragments engages Toll-like receptor 4 (TLR4) and its adaptor protein MyD88, which are thought to play critical roles in the development of neuronal pathology through mechanisms that remain incompletely understood. For example, in MPS IIIB mice, attenuation of TLR4 alone or in combination with MyD88 reduced neuroinflammation but did not abolish secondary neuronal damage [37]. Studying the cell-type-specific pathology in the animal models of neurological LSD remains challenging due to complexity of the CNS and inherent limitations of *in vivo* modeling. In this context, co-cultures of patient-derived neuronal and glial cells provide a powerful platform to dissect both cell-autonomous and non-cell-autonomous pathological mechanisms. Here, for the first time, we demonstrate that iMGL derived from MPS IIIC iPSCs not only recapitulate *bona fide* microglial phenotype but also exhibit disease-specific features such as lysosomal enlargement and increased ILB4 levels, findings consistent with prior observations in MPS IIIC mouse models [38]. Furthermore, our results revealed that MPS IIIC microglia acquire an activated phenotype and contain lysosomal puncta positive for the synaptic marker VGLUT1 and the dendritic marker MAP2, consistent with excessive synaptic pruning and degradation of neuronal projections observed in co-cultured MPS IIIC iCN.

In contrast, MPS IIIC iMGL overexpressing HGSNAT following gene correction and producing HGSNAT^+^ EV effectively rescued most disease-associated phenotypes in co-cultured MPS IIIC iCN. This intervention led to complete normalization of lysosomal LAMP2^+^ areas, clearance of the primary HS^+^ storage, secondary G_M2_ accumulation, and restoration of synaptic protein levels, effects that were not achieved by short-term treatment of neurons with purified HGSNAT^+^ EV. These findings highlight the importance of sustained exposure for complete phenotypic correction.

Notably, a similar rescue of lysosomal enlargement and synaptic marker deficits was also achieved when MPS IIIC iCN were treated with conditioned media from iMGL overexpressing HGSNAT-EGFP, depleted of detached cells and cell debris, but retaining EV. Collectively, these results provide compelling evidence that iMGL overexpressing HGSNAT-EGFP mediate cross-correction of co-cultured MPS IIIC iCN via secretion of HGSNAT^+^ EVs rather than through the direct transfer of intact lysosomes via “tunneling nanotubes”, a mechanism previously reported for fibroblasts deficient in the lysosomal cystine transporter co-cultured with normal HSPC-derived macrophages [39].

A recent study also demonstrated correction of GAG storage in MPS IIIC fibroblasts treated with EV produced by cells transduced with an rAAV-hHGSNAT^EV^ vector containing an EV-mRNA-packaging signal in the 3’UTR, designed to promote inclusion of *HGSNAT* mRNA into EV [40]. The authors attributed the observed therapeutic effect to translation of *HGSNAT* mRNA delivered by EV, resulting in production of the functional enzyme in the recipient cells [40]. However, based on the results presented in this study, it cannot be excluded that the EV secreted by AAV-transduced cells overexpressing HGSNAT may also carry functional HGSNAT enzyme and deliver it directly to the recipient cells.

Overall, our work provides a robust proof-of-concept for cross-correction of CNS and peripheral cells using EV produced by cells derived from transplanted, gene-corrected HSPC. Importantly, CNS pathology in MPS III and other neurologic LSD is characterized by chronic microglial and astrocytic activation accompanied by elevated levels of pro-inflammatory cytokines and chemokines [37, 41]. This inflammatory milieu promotes monocyte and macrophage infiltration into the brain parenchyma, further exacerbating neurodegeneration [5, 38]. Transplantation of wild-type HSPC into MPS IIIC *Hgsnat^P304L^* knock-in mice has been shown to reduce microglial activation (as indicated by reduced CD68^+^ cell density) and cytokine expression (decreased *IL-1β* expression), resulting in partial behavioral improvements. However, the therapy failed to correct neuronal HS storage or memory deficits [38]. In this context, gene-targeted microglia overexpressing HGSNAT may exert a more comprehensive therapeutic effect by mitigating neuroinflammation and enabling cross-correction of neurons and other CNS cells through the release of enzyme-loaded EV. Compared with other vector-based approaches, LV-mediated EV production in human cells supports long-term, potentially life-long expression of therapeutic enzymes, while posing a lower risk of immune complications and absence of adverse effects despite supraphysiologic enzyme levels in myeloid lineages and plasma [19, 20]. Our experiments, likewise, revealed no detectable detrimental effects in cultured control neurons treated with either conditioned iMGL media or purified HGSNAT^+^ EV, although additional *in vivo* studies are warranted to confirm safety. Despite the potential of EV to cross the BBB, their efficiency in CNS penetration remains limited. Moreover, endothelial and perivascular abnormalities previously reported in MPS III may present additional challenges for effective blood-to-CNS delivery[42].

One drawback of this study is the limited functional characterization of MPS IIIC neurons treated with HGSNAT^+^ EV or cocultured with iMGL overexpressing the enzyme. In particular, detailed analysis of their electrophysiological activity, synaptic ultrastructure, and gene or protein expression profiles, similar to those we previously performed for untreated MPS III human iCN and mouse primary hippocampal neurons [33], are required to further confirm the therapeutic efficacy and assess potential adverse effects. Future studies should also focus on optimizing the lentiviral transduction protocols during the iHPC stage and/or employ flow cytometric enrichment of HGSNAT-EGFP^+^ microglial progenitors prior to differentiation. This could lead to higher therapeutic potency once these cells mature into microglia. Currently, there are no established methods to maintain transduced iHPC lines for long-term culture, as this stage is inherently transient and typically used for rapid downstream differentiation or transplantation experiments [26, 43, 44]. Nevertheless, even a mixed population of transduced microglia was sufficient to significantly ameliorate MPS IIIC neuronal pathology in co-cultured iCN. A comprehensive characterization of microglial subtypes before and after transduction using IF, flow cytometry, and single-cell transcriptomic profiling will further elucidate microglial responses to disease and HGSNAT overexpression.

In conclusion, this study demonstrates the feasibility and efficacy of producing HGSNAT-loaded EV using either human HCMEC/D3 cells or iPSC-derived microglia transduced with LV HGSNAT-EGFP constructs for delivery of the therapeutic enzyme to other cell types, including neurons. Co-culturing MPS IIIC iCN with iMGL engineered to overexpress HGSNAT resulted in a complete phenotypic rescue, underscoring the translational potential of EV-mediated enzyme delivery to mitigate the CNS pathology in MPS IIIC. Further studies should aim to extend this approach to other lysosomal storage diseases, and therapeutic enzymes and proteins, as well as to explore the potential of combining EV-mediated enzyme delivery with complementary therapeutic modalities.

## Materials and Methods

### Availability of data and materials

The data, analytic methods, and study materials will be made available to other researchers for purposes of reproducing the results or replicating the procedure.

### Study approval

Generation and analysis of iPSC-derived cortical neurons of MPS III patients was approved by the Ethical research board of CHU Ste-Justine (approval number 2022-3817).

### Enzyme activity assays

To measure HGSNAT activity, 5 µl aliquot of cell homogenate was combined with 5 µl of McIlvain Buffer (pH 5.5), 5 µl of 3 mM 4-methylumbelliferyl-β-D-glucosaminide (Moscerdam), 5 µl of 5 mM acetyl-coenzyme A and 5 µl of H_2_O and the reaction mixture was incubated for 3 h at 37°C. The reaction was stopped by adding 975 µl of 0.4 M glycine buffer (pH 10.4), and fluorescence was measured using a ClarioStar plate reader (BMG Labtech). Blanks for each sample were incubated without the homogenates for the 3 h, and the homogenate added after the glycine buffer [25].

### hCMEC/D3 cell cultures and LV transduction

Human immortalized cerebral microvascular endothelial cells (hCMEC/D3) [29] were the kind gift of Dr. Pierre Olivier Couraud (Cochin Institute, Université Paris Descartes). Cells were cultured at 37°C in a humidified incubator at 5% CO₂ and 95% O₂, using EBM-2 basal medium (Lonza, Walkersville, MD, USA) supplemented with 25% (v/v) of a SingleQuot kit (Lonza) and 2% fetal bovine serum (FBS). Cultures were maintained in flasks pre-coated with 100 µg/mL of rat tail collagen type I (BD Canada, Mississauga, ON), diluted in 20 mM acetic acid. Cells from passages 30 to 34 were used for experiments.

### Lentiviral transduction

Lentivirus CD68-HGSNAT-EGFP was produced using the 293SF-PAC-LV-IIIb-3D4 packaging cell line [45]. Briefly, the packaging cells were transfected with a mixture of PEI and the required plasmid. Four hours post-transfection (p.t.), cumate and coumermycin were added to induce the production of lentiviral elements. The following day, the cells were concentrated fourfold in medium containing cumate, coumermycin, and sodium butyrate. Three days p.t., the culture supernatant was collected, concentrated approximately 600-fold by ultracentrifugation, and titrated using a gene transfer assay. Lentivirus CMV-HGSNAT-EGFP used for expression of HGSNAT-EGFP under the control of the CMV promoter was produced by co-transfecting HEK293T cells with pLenti-HGSNAT-GFP [25], REV (Plasmid #12253; Addgene), pMDL (gag-pol; Plasmid #12251; Addgene), and VSV-G (Plasmid #12259; Addgene) using Lipofectamine 2000 (Invitrogen, 11668019). Culture supernatants containing lentivirus were collected 30 h post-transfection and used to transduce hCMEC/D3 cells using standard protocol in the presence of polybrene (8 mg/ml). After 3 days, cells were treated for 5 days with the media containing 2 µM puromycin to select transduced cells and then EGFP-positive cells were purified by cells sorting. Transduction was confirmed by observing the Turbo-GFP fluorescent signals using a ZOE Fluorescent Cell Imager.

### Extracellular Vesicles (EV) Isolation

hCMEC/D3 cells overexpressing HGSNAT-GFP were grown in T-75 flasks under serum-free conditions as previously described [31] to avoid contamination from serum-derived EV from FBS and minimize non-specific binding of serum components to hCMEC/D3-derived EV. Conditioned media (∼100 mL) was collected every 24 hours over three days and pooled. The media was further subjected to differential centrifugation to isolate purified EVs as previously described [31]. Briefly, the media was centrifuged at 300 x *g* for 10 min to remove any intact cells. The supernatant was further centrifuged at 10,000 x *g* for 30 min to pellet macrovesicles. The media was then transferred to an ultracentrifuge tube and centrifuged at 100,000 x *g* for 70 min using an Optima TLX ultracentrifuge equipped with 60 Ti rotor (Beckman Coulter, Mississauga) to pellet EV. The supernatant was moved to a new tube and centrifugation repeated. The pellets were resuspended in PBS combined and centrifuged at 100,000 x g for 70 min. The pelleted EV were resuspended in DMEM media (without sodium pyruvate, with 1.5 g/L sodium bicarbonate, with 4.5 g/L glucose, with L-glutamine) and stored at 4°C to be used in subsequent experiments.

### Characterization of EV

EV were analyzed using the ZetaView PMX-420 nanoparticle Tracking analyzer (Particle Metrix, Germany) equipped with the software version 8.05.11 SP4. After an initial calibration using ZetaView 100 nm alignment suspension (diluted 1:250,000 in milliQ filtered water), the instrument was thoroughly rinsed with PBS and 1 ml of EV sample, containing 1x10^4^ - 1x10^6^ particles, was injected. Once the sensitivity value, particles/frame and the particle drift was within acceptable ranges, video acquisition was run under the following settings: Sensitivity 85, Shutter Speed 40, Frame Rate (fps) 30, Resolution Highest, Camera Gain 770, Positions Measured 11, Minimum Brightness 15, Minimum Size (pixels) 10, Maximum Size (pixels) 500.

### Simple Western blot analysis

Simple Western blot technology (ProteinSimple Wes) was used for EV characterization. This fully automated capillary-based immunoblotting system combines protein separation by size or charge with sensitive chemiluminescence or fluorescence immunodetection using conventional primary and conjugated secondary antibodies. Approximately, 0.5 µg of EV protein (3.28 x 10^5^ EV particles) was loaded per capillary and analyzed under reducing and denaturing conditions using a 12-230 kDa separation module (ProteinSimple; Catalogue#SM-W001). Primary antibodies included anti-CD9 (84 µg/ml; Cell Signaling 13174), anti-CD63 (0.5 mg/ml; Boster PB9250), anti-TSG101 (0.5 mg/ml; Boster RP1078) and anti-Flotillin-2 (C42A3) 19 µg/m (Cell Signalling 3436). Anti-Calnexin antibody (1 mg/ml; (Abcam ab22595) was used to confirm the purity of EV preparation. HGSNAT-GFP fusion protein in the stock EVs (2.9 x 10^9^ EV particles) was detected using anti-GFP at 1:100 dilution (Abcam ab290). Detection of primary antibodies was performed with anti-rabbit IgG antibodies (ProteinSimple; Catalogue#DM-001).

### Immunocytochemistry

Cultured hCMEC/D3 cells, fibroblasts, iCN at DIV28 or DIV34, and iMGL at DIV28 or DIV34 were fixed using a 4% paraformaldehyde in PBS, pH 7.4, for 15 min. The cells were permeabilized with 0.1% Triton-X100 in PBS for 5 min, and non-specific binding sites were blocked with 5% BSA (Wisent) in PBS for 1 h. The cells were then, incubated overnight at 4°C with primary antibodies in 1% BSA in PBS (see Table 1 for the source of antibodies and their dilutions). On the following day, the cells were washed 3 times with PBS and labelled with appropriate Alexa Fluor 488-, Alexa Fluor-555- or Alexa Fluor 633-conjugated anti-IgG antibodies (1:1000, all from Thermo Fisher Scientific) for 1 h in 1% BSA in PBS at room temperature, washed 2 times using PBS, and once with dH_2_0. Coverslips were mounted on glass slides using ProLong Gold mounting medium, containing 4′,6-diamidino-2-phenylindole (DAPI; Invitrogen, Cat # P36935), and analyzed by a Leica SP8-DSL confocal microscopes (× 40 and x 63 glycerol immersion objectives, N.A. 1.4). Images were processed with Leica Application Suite X (LAS-X) software and Photoshop 2021 (Adobe) and quantified using Fiji-ImageJ 1.50i software (National Institutes of Health, Bethesda, MD, USA). Analysis of images was performed with summation of 10 z-stacks separated by 0.5 µm. Neuronal soma or axon areas were identified by NEUN, or MAP2 labeling. To measure LAMP2^+^ area per neuron total LAMP2+ areas were normalised with iCN NEUN^+^ area. LAMP2^+^ and ILB4^+^ areas were normalized with IBA1+ areas for iMGL. Synaptic puncta were counted along 10-30 µm segments of neuronal projections at a <10 µm distance from the soma. Colocalization was estimated using the Person’s correlation coefficient. Quantification was blinded and performed for 3 independent experiments.

**Table 1:**
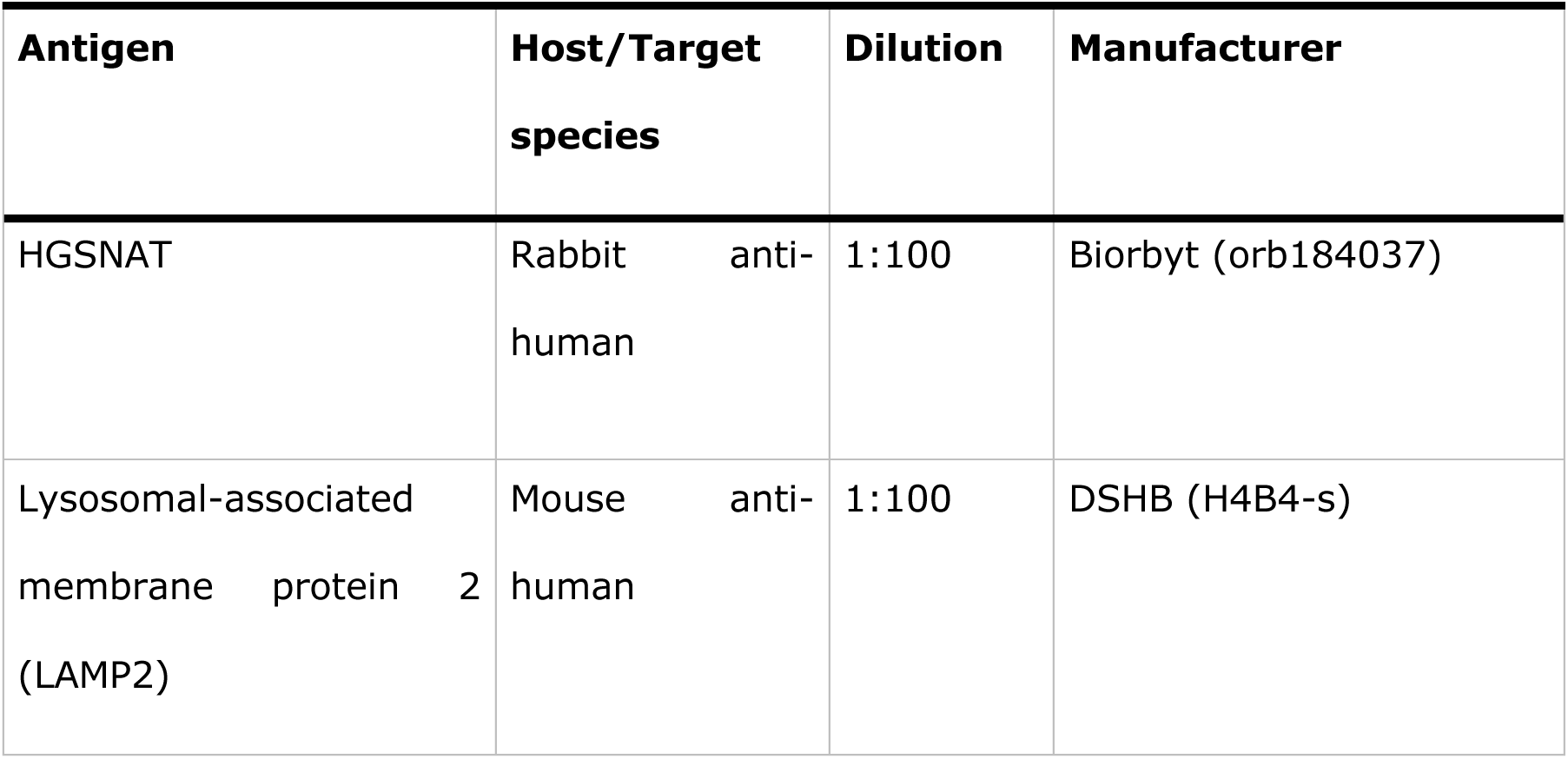
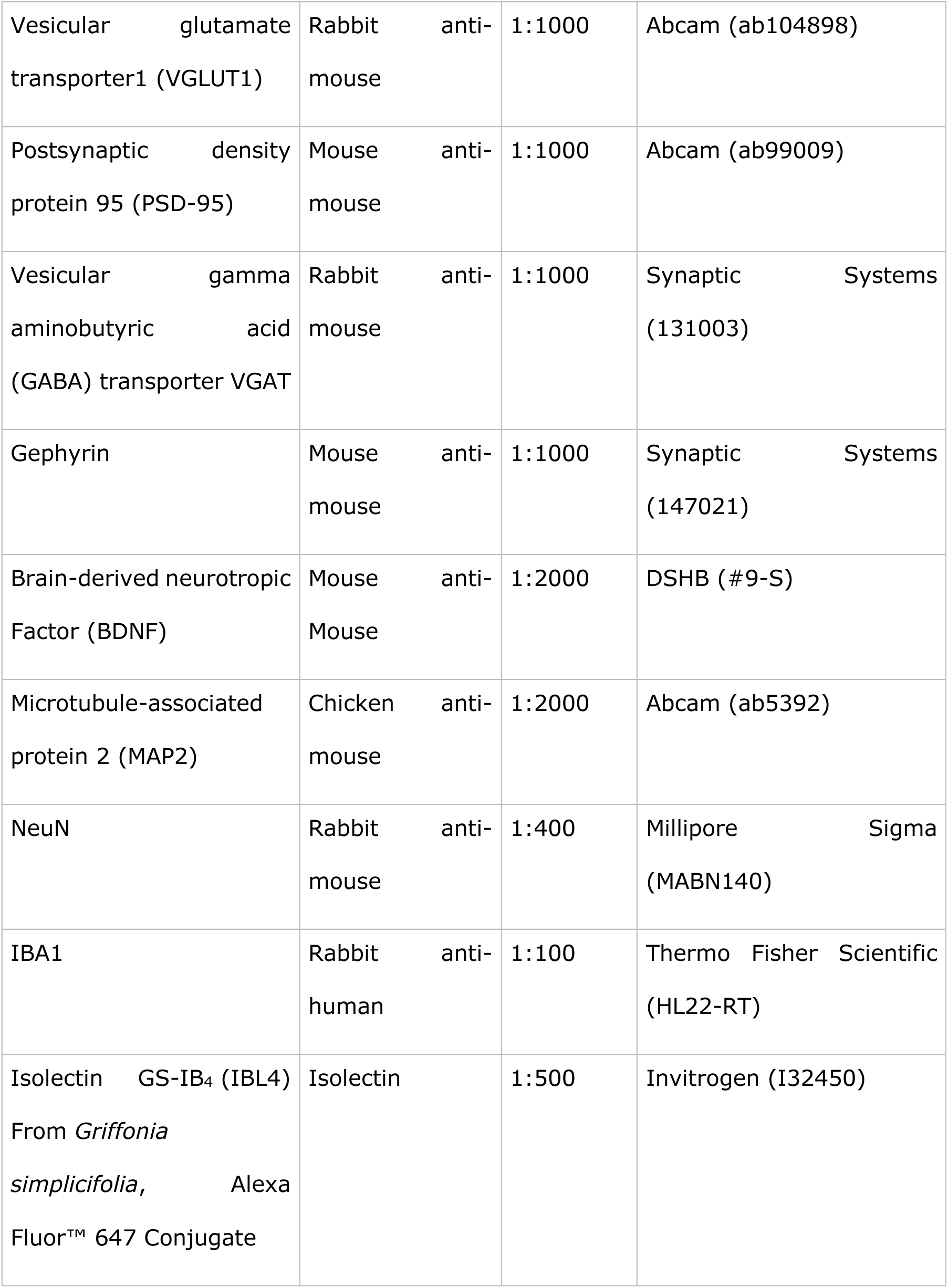
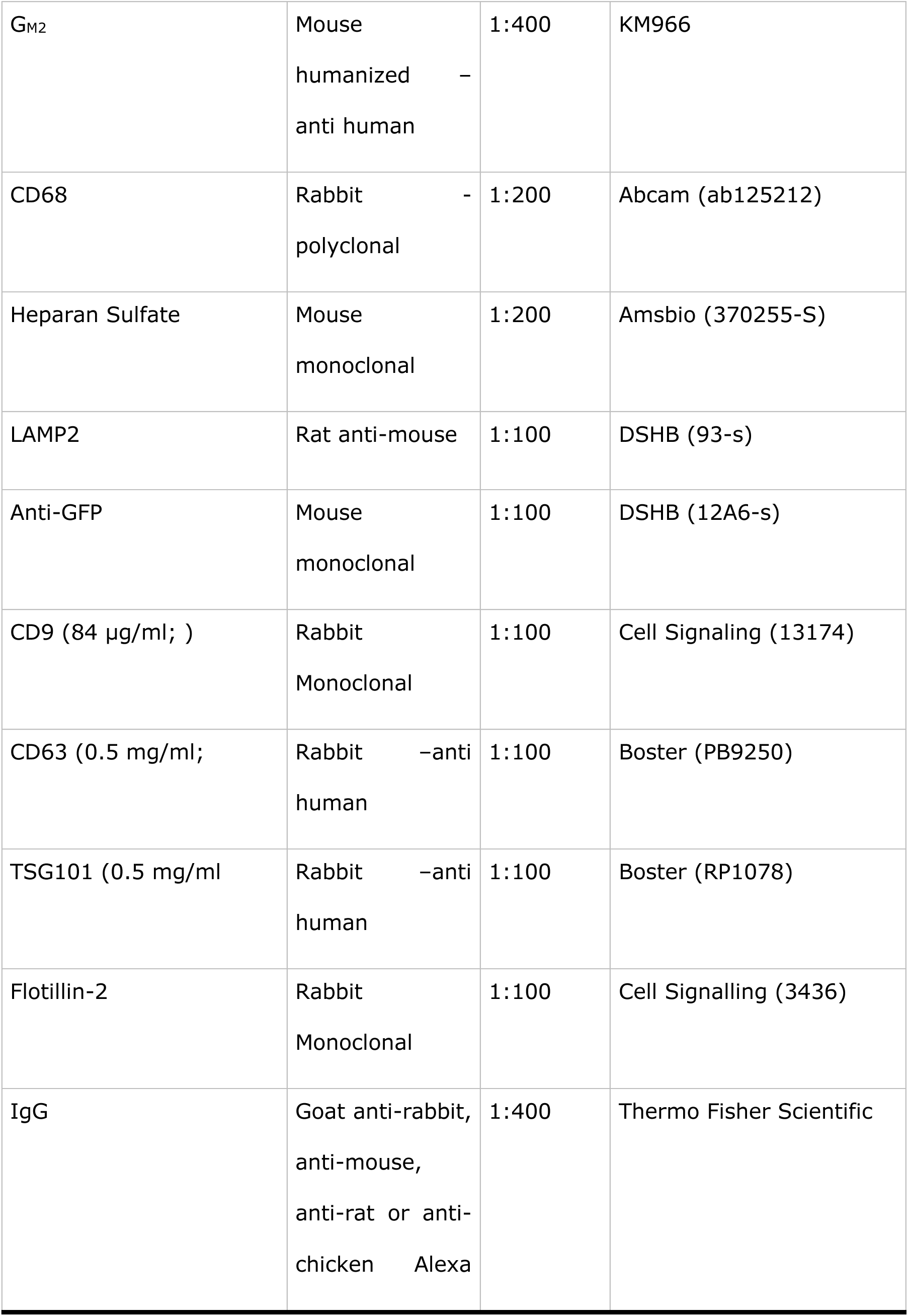
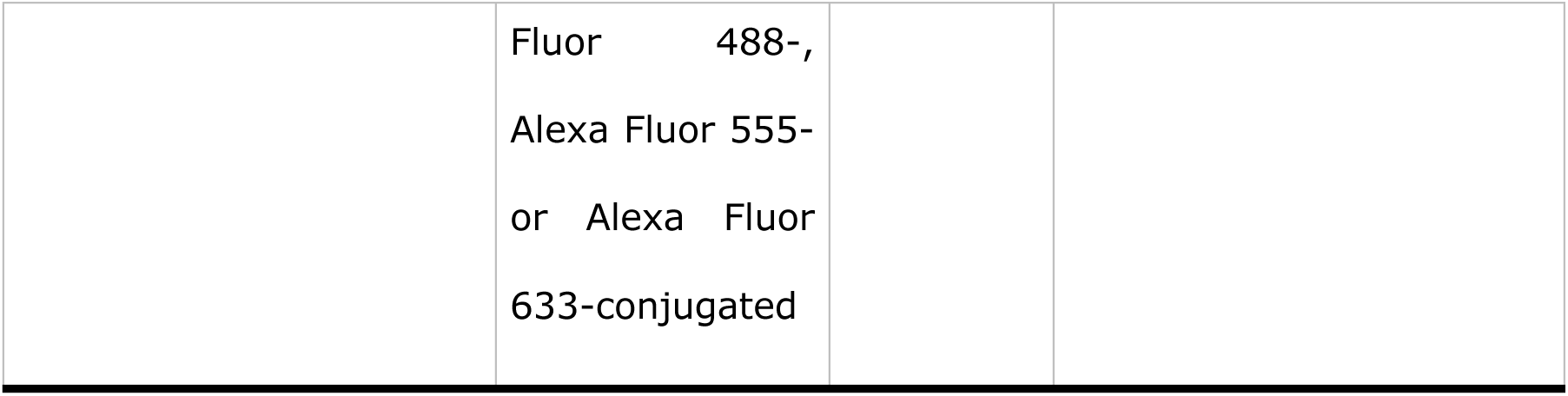
Antibodies used in the study and their working concentrations

### Generation and analysis of iPSC-derived neuronal cortical neurons (iCN)

Generation of iPSC lines was previously described [32], (Moore, Dubot et al., bioRxiv 2026.02.20.707013). In the current study, we used two control lines (CNT-1, and CNT-2) and one MPS IIIC patient fibroblast line. iPSC were expanded and maintained on six-well plates coated with Matrigel^TM^ mTeSR™ Plus medium at 37°C, in 5% CO_2_/5% O_2_ atmosphere following the medium manufacturer’s protocol [46]. At 60-80% confluency the cells were passaged using Gentile Cell Dissociation reagent (GCDR, StemCell Technologies^TM^) and plated in mTeSR™ Plus medium containing 10 µM RI (Y27632 ROCK inhibitor, Selleckchem). The following day, the medium was replaced by fresh mTeSR™ Plus medium without RI.

iPSCs were differentiated into cortical forebrain committed neural precursor cells (NPCs) by dual SMAD inhibition, as described [47] by passaging iPSCs to Matrigel^®^ coated dishes with subsequent passaging to poly-L-ornithine (PO)/laminin coated dishes. NPC induction was performed in a monolayer with the cortical neuronal induction media essentially as described [47] but with FGF-8 used instead of FGFb-2. Eighty percent of media was changed every 2 days. After induction for 3 weeks NPCs were differentiated into iCNs neurons as previously described [48]. First, NPCs were passage into PO/laminin coated plates in a 1/1 mixture of DMEM/F-12, GlutaMAX™/Neurobasal™ (NB, Gibco^TM^) media containing B27, N2, NEAA, BDNF, GDNF, Laminin, dbCAMP, Compound E and TGF-B3 containing 2 µM RI. The following day, media was changed for a 100% NB media containing the above components. Neurons were then cultured for 4 weeks (DIV28) until fully differentiated and matured.

### Treatment of iCN cultures with HGSNAT^+^ EV

HGSNAT^+^ EV isolated from the medium conditioned by hCMEC/D3 overexpressing HGSNAT-EGFP and resuspended in fresh NB media were added to mature iCN cultures at DIV 25 during the regular media change at a dose of 1.30 10^6^ particles per well. After 54 h in culture, the cells were fixed and analysed.

### Generation and analysis of iPSC-derived microglia (iMGL)

iPSC-derived hematopoietic progenitor cells (iHPC) were generated using the STEMdiff^TM^ Hematopoietic kit (STEMCELL Technologies) as previously described [26, 49]. Small iPSC colonies were detached and plated into 6-well plates coated with Matrigel^TM^. 24 h after passaging the cells were induced in STEMdiff^TM^ Hematopoietic media for 16 days. Floating iHPC were collected on days 12, 14, and 16 and counted before conversion into microglia or cryopreserved using BAMBANKER® (FujiFilm, Wako Chemicals). iHPC were matured into iMGL for 28 days using the protocol established by Dorion et al. [26]. Briefly, iHPC were replated at a density of 1-2 x10^6^ cells/well on fresh Matrigel^TM^ coated 6-well plates and cultured in the maturation media (2 ml/well) of MEM alpha (Gibco^TM^) containing B27, Insulin-transferrin-selenite, Glutamax, and three cytokines IL-34, TGFb, M-CSF. Fresh maturation media (1 ml/well) was added three times after 3, 7 and 10 days of culture. On the 14^th^ day of culture, the conditioned media was collected, centrifuged at 300 *g* for 5 min, mixed with fresh media at 1:1 ratio and added back to the cells. The procedure was repeated by adding fresh media until DIV 25, where along with the spun conditioned media two additional cytokines GM-CSF and CX3CL1 were added to the freshly added maturation media. The procedure continued until DIV 28, when the cells reached maturity and could be used for the downstream experiments.

### Transduction of iHPC with LV-CD68-HGSNAT-EGFP or LV-CMV-EGFP

iHPCs on day 12 were transduced with the LV CD68-HGSNAT-EGFP (1.89 x 10^9^ transduction units (TU)/mL) or LV CMV-EGFP (8.3 × 10^8^ TU/ml) generously provided by Dr. Jeffrey A. Medin (Medical College of Wisconsin, Milwaukee, USA). To maximize transduction efficiency, the virus was added twice with 24 h interval at the multiplicity of infection (MOI) 30 in the presence of polybrene (Millipore Sigma) at 1:1000 dilution.

### iMGL/iCN co-cultures

Replating of DIV 25 or 28 iMGL followed the replating protocol for iMGL cells [26] with slight modifications. After the iMGL were collected by centrifugation, the cells were resuspended in 1:1 mixture of conditioned and fresh differentiation medium (with all five cytokines). After counting, iMGL were re-plated on Matrigel^TM^ coated coverslips in 24-well plates at a density of 5-7.5 x 10^4^ cells/well to mature as monocultures until DIV 28. To establish combined iMGL/iCN co-cultures, iMGL were seeded over iCN cultured on PO/laminin-coated coverslips in 24-well plates at a 2:1 ratio of iCNs to iMGL (∼10^5^ iCNs to 5 10^4^ iMGL) and cultured until DIV 35. in a 50:50 mixture of iCN and iMGL media.

### Phagocytic assay with pHrodo™

To test iMGL activation and immune response, the iMGL at DIV 28 were treated with LPS at a final concentration of 100 ng/ml for 24 h. On the following day, cells were fixed with 4% PFA before analysis for IF microscopy. Analysis of phagocytosis using pHrodo™ Green-labelled *Escherichia coli* (*E. coli*), Bioparticles™ from Thermo Fisher Scientific followed manufactures protocol modified for 24-well plates. The powdered pHrodo™ was dissolved in PBS at the concentration of 1 mg/mL and the solution transferred into a 15 ml tube, vortexed and sonicated on ice for 10 min. Twenty μl of the pHrodo™ suspension was added to 500 µl of media in each well of the 24-well plate, followed by a gentle rotation, and cells were incubated for 3 h at 37°C. Cells were then washed 6 times for 10 min each with PBS before fixation and analysis.

### iPSC lines of MPS IIIC patient and normal controls used in this study

iPSC lines derived from two healthy controls (CNT-1 and CNT-2) and MPS IIIC patient (MPS IIIC-1) used in this study were previously described in [32] and (Moore, Dubot et al., bioRxiv 2026.02.20.707013).

### Collection of iMGL-conditioned media and treatment of iCN cultures

Conditioned media was collected from the 6-well plates of iMGLs at DIV 28 and centrifuged as described to remove floating cells, cell debris, apoptotic bodies and large EVs (>1,000 μm) [35]. Briefly, media was centrifuged at 300 x *g* for 10 min to remove residual cells, then media was transferred to a new tube and centrifuged at 2,000 x *g* for 10 min. Media was then transferred to a new tube and centrifuged at 10,000 x *g* for 10 min. Conditioned media from CNT and LV CD68-HGSNAT-EGFP-transduced MPS IIIC iMGL were added in a 1:1 (v/v) ratio to the NB media of CNT and MPS IIIC iCN cells in 24 well-plates for 56 h before fixation and analysis.

## Acknowledgments

The authors thank Tara Khayer and Valero Piscopo for helping advice, Dr. Elke Küster-Schöck and the Plateforme d’Imagerie Microscopique (PIM – CHU Sainte Justine) for the help with confocal microscopy, Dr. Mila Ashmarina for critically reading the manuscript and Ewa Baumann for her help with the Wes data. The authors are also grateful to Richard Gingras for the assistance with plasmid production and purification.

## Author contributions

Author contributions: T.M., M.T., X.P., E.B., M.H., D.L.M., M.F., A.B., M.E., J.K.S., M.R., C.C. and A.J. conducted experiments and acquired data; T.M., M.T., X.P., E.B., M.H., D.L.M., A.B. M.E., J.K.S., A.J. and A.V.P. analysed data; F.A., T.D., M.E., J.K.S., A.J. provided essential resources; T.M., M.T., X.P., J.K.S., A.J. and A.V.P. designed the experiments; T.M. and A.V.P and wrote the manuscript (first draft); T.M., J.K.S., A.J., and A.V.P. edited the manuscript. All authors read and approved the final version of the manuscript.

