## Supplemental figure for "Extracellular vesicles derived from cells overexpressing HGSNAT rescue defects in Mucopolysaccharidosis IIIC neurons"

### Supplementary Materials

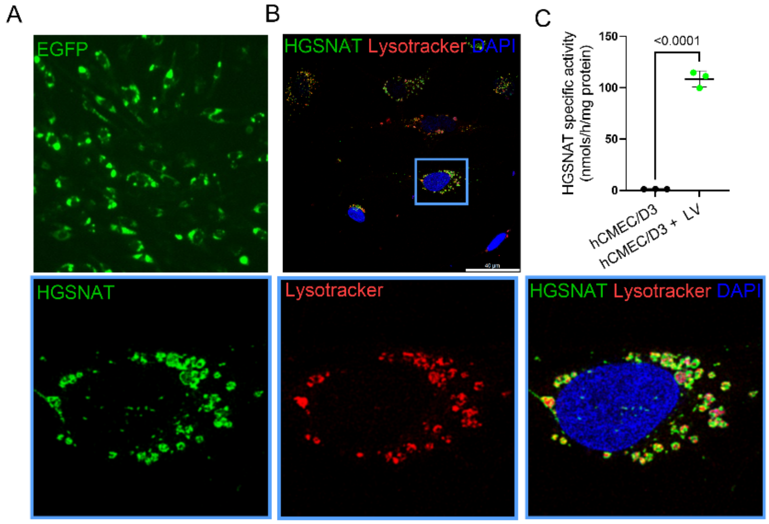


**Figure S1: hCMEC/D3 cells transduced with LV CMV-HGSNAT-EGFP virus reveal localization of HGSAT-GFP in the lysosomes and supraphysiological levels of HGSNAT activity.** (**A**) LV-CMV-HGSNAT-EGFP transduced hCMEC/D3 cells show the EGFP$^{+}$ (green) puncta post-sorting. (**B**) HGSNAT-EGFP$^{+}$ perinuclear puncta show colocalization with the lysosomal marker LysoTracker (red). DAPI (blue) was used to label nuclei. The scale bars equal 40 µm. Inserts show enlarged images of selected hCMEC/D3 cell (blue rectangle). (**C**) Post-sorted LV-CMV-HGSNAT-EGFP transduced hCMEC/D3 cells reveal supraphysiological levels of HGSNAT activity.

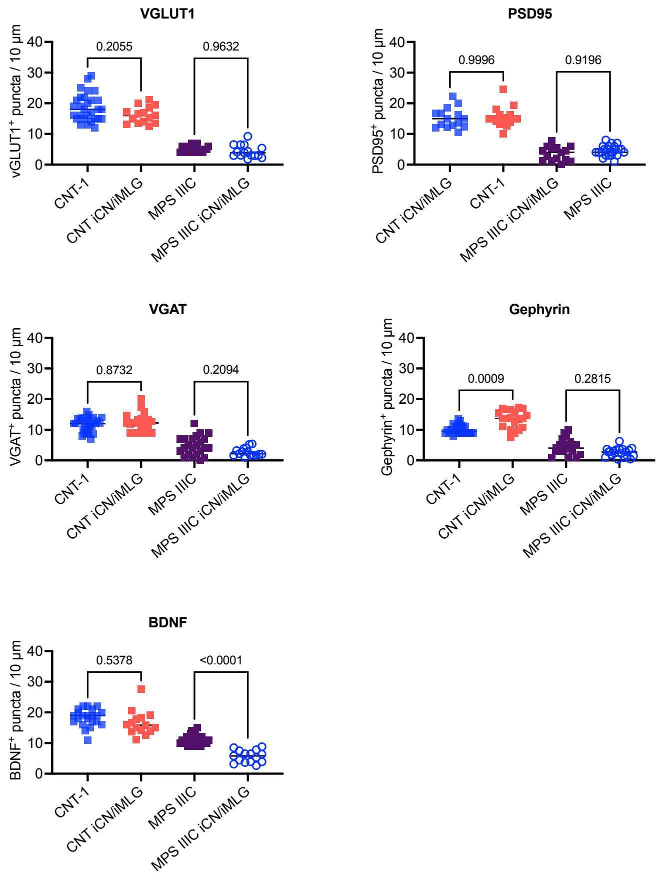


**Figure S2: Comparison of synaptic proteins and BDNF puncta levels in iCN monocultures and iCN/iMGL co-cultures.**

Quantification of VGLU1$^+$, PSD95$^+$, VGAT$^+$, Gephyrin$^+$, and BDNF$^+$ puncta along 10-μm segments of neuronal projections was performed with ImageJ. Levels of BDNF$^+$ puncta are reduced in MPS IIIC iCN/iMGL co-cultures compared to MPS IIIC iCN monocultures. Graphs show individual results, means and SD from three independent cultures (≥15 cells in each experiment). *P*-values were calculated using nested one-way ANOVA and Tukey post hoc test.

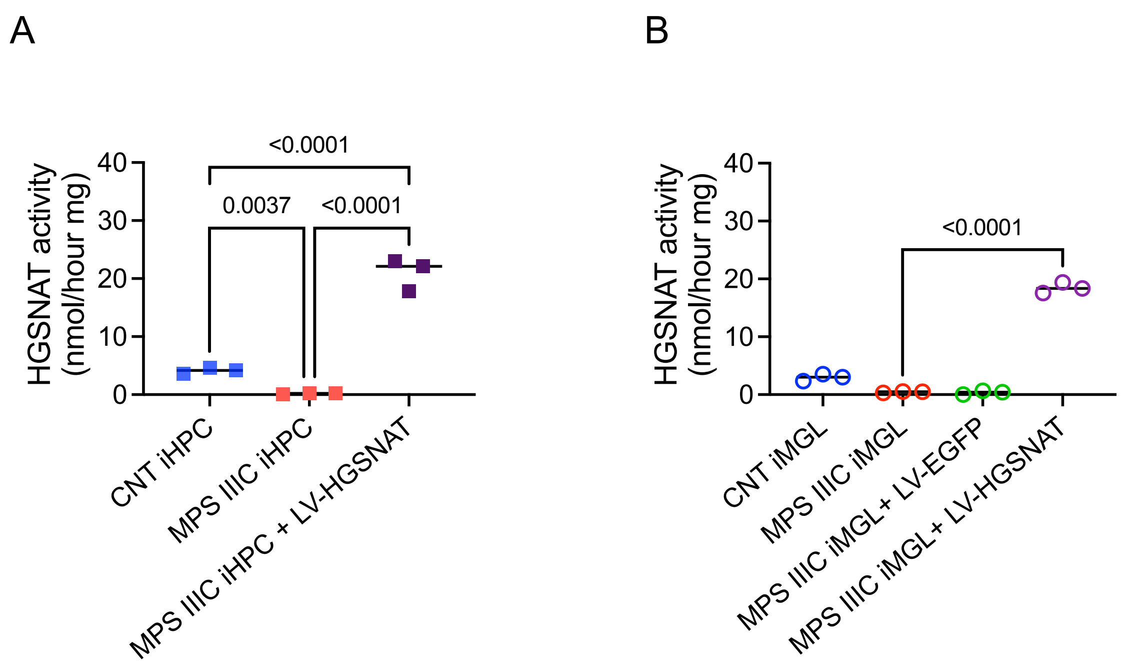


**Figure S3: HGSNAT activity in MPS IIIC iHPC and iMGL transduced with LV CD68-HGSNAT-EGFP at MOI 30 is increased to supraphysiologic levels.**

(**A**) HGSNAT activity was measured with the fluorogenic substrate in the cells pellets of normal control iHPC (CNT) and MPS IIIC iHPC transduced or not with the LV CD68-HGSNAT-EGFP vector at MOI 30. iPSC-derived iHPC of MPS IIIC patient show characteristic primarily enzymatic deficit of HGSNAT activity. The HGSNAT activity in MPS IIIC iHPC transduced with the LV CD68-HGSNAT-EGFP vector is increased ~8-fold compared to normal control cells. (**B**) HGSNAT activity was measured in the cell pellets from DIV 28 iMGL derived from normal control iHPC (CNT) and MPS IIIC iHPC transduced or not with the LV CD68-HGSNAT-EGFP vector at MOI 30. The HGSNAT activity in MPS IIIC iMGL is reduced to <1% of the activity in the control cells. The MPS IIIC iMGL transduced with the LV CD68-HGSNAT-EGFP vector show supraphysiological levels of HGSNAT activity. Graphs show individual results, means and SD, for three individual cultures. Duplicate measurements were performed for each culture. *P*-values were calculated using one-way ANOVA with Tukey post hoc test.
